# Large Fatty Acid-derived Aβ42 oligomers Form Ring-like Assemblies

**DOI:** 10.1101/390567

**Authors:** Wenhui Xi, Dexter N. Dean, Kelli A. Stockmal, Sarah E. Morgan, Ulrich H.E. Hansmann, Vijayaraghavan Rangachari

## Abstract

As the primary toxic species in the etiology of Alzheimer disease (AD) are low molecular weight oligomers of Aβ, it is crucial to understand the structure of Aβ oligomers for gaining molecular insights into AD pathology. We have earlier demonstrated that in the presence of fatty acids Aβ42 peptides assemble as 12-24mer oligomers. These <u>L</u>arge <u>F</u>atty <u>A</u>cid-derived <u>O</u>ligomers (LFAOs) exist predominantly as 12mers at low, and as 24mers at high concentrations. The 12mers are more neurotoxic than the 24mers and undergo self-replication, while the latter propagate to morphologically distinct fibrils with succinct pathological consequences. In order to glean into their functional differences and similarities, we have determined their structures in greater detail by combining molecular dynamic simulations with biophysical measurements. We conjecture that the LFAO are made of Aβ units in an S-shaped conformation, with the 12mers forming a double-layered hexamer ring (6 × 2) while the structure of 24mers is a double-layered dodecamer ring (12 × 2). A closer inspection of the (6 × 2) and (12 × 2) structures reveals a concentration and pH dependent molecular reorganization in the assembly of 12 to 24mers, that seems to be the underlying mechanism for the observed biophysical and cellular properties of LFAOs.

## I. INTRODUCTION

One of the hallmarks of Alzheimer disease (AD) pathology is the deposition of amyloid-β (Aβ) peptide fibrillar aggregates as plaques in brains of patients. The neuronal loss, however, seems to be triggered primarily by low-molecular weight (LMW) oligomers that are formed earlier than the high-molecular weight fibrils during the aggregation process ^1^. Therefore, there is a growing interest in isolating LMW oligomers and deriving their atomistic structure and dynamics. However, the transient nature and heterogeneity of the oligomers makes their isolation and characterization, either from endogenous or exogenous sources, difficult. As a consequence, deriving structural models, as needed for understanding their toxicity mechanism and mode of propagation, poses a challenge.

We have developed a method for generating distinct Aβ42 12-24mer assemblies called, large fatty acid-derived oligomers (LFAOs) ^2-7^ as their generation is catalyzed by saturated fatty acids. At higher concentrations, LFAOs convert from the 12mer species to more disperse distribution of 12-24mer oligomers ^2^. This concentration-dependent transition is significant because the 12mers self-replicate in the presence of monomers, and are more apoptotic than the 24mers ^2^. On the other hand, the 24mers faithfully propagate towards morphologically-unique fibrils and induce acute cerebral amyloid angiopathy (CAA) in transgenic mice ^3^. Therefore, it is imperative to obtain atomistic insights into the differences between the 12mer and 24mer LFAOs, as well as the transition from one form to the other, in order to better understand their unique properties. Unfortunately, due to some of the aforementioned reasons, the structures of these LFAOs have so far not been resolved.

Guided by computational investigations of the Hansmann laboratory, we present in this paper structural models for 12mer and 24mer LFAOs, and a mechanism for the transition from one to the other assembly. Derivation of these models relies on the computational techniques developed in the Hansmann lab. These methods have been already used in previous work to study the various polymorphs seen in amyloids ^8-11^, and to probe the factors that modulate conformational switching in amyloids ^11^. Combining computational investigations with novel biophysical experiments, we conclude that the 12mers prefer to form two-layered rings, each ring a hexamer (6 × 2), while 24mers transition to another species of two-layered assemblies, here each ring a dodecamer (12 × 2). We also eliminate the possibility of a single dodecamer ring (12 × 1) structure for 12mers, and of four stacked hexamer rings (6 × 4) as the structure for 24mers. Our models allow us to explain the experimentally observed conformational-dynamics of LFAOs, to identify the key residues involved in conformational switching, and the provide hints at the structural basis for the different pathogeny of LFAO 12mers and 24mers.

## II. METHODS

### A. Materials

Lyophilized stocks of synthetic Aβ 1-42 WT peptide were procured from the Yale School of Medicine peptide synthesis facility (New Haven, CT). C12:0 NEFA was purchased from NuCheck Prep, Inc. (Elysian, MN), while ANS was purchased from Sigma-Aldrich Corp. (St. Louis, MO). All other buffers, reagents, and consumables were procured from Thermo Fisher Scientific, Inc. (Waltham, MA).

### B. Molecular dynamics simulation

The oligomers models presented in this article are based on previous work where we constructed a series of N-fold ring-like Aβ42 oligomer models ^8^, including six-fold and twelve-fold models of Aβ(11-42), that are characterized by S-shaped chain configurations forming three β-strands liked by two turn regions. The chains in a ring are kept together by hydrophobic contacts in the region of residues 20-28 and an inter-chain salt bridge K16-D23. These models are the starting point in our construction of the (6 × 2), (6 × 4) and (12 × 2) models described later.

Our simulations rely on the software package GROMACS 5.1.5 ^12^ end employ the AMBER ff99SB-ildn ^13^ force field for proteins and TIP3P water ^14^ as solvent, a choice also employed by us in our previous work ^15^. The temperature of 300K and a pressure of 1 bar are controlled by v-rescale thermostat ^16^ and Parrinello-Rahman barostat ^17^. The bond-lengths are restrained by the LINCE algorithm ^18^ and SETTLE algorithm ^19^ allowing us to use a time step of 2 fs for integration. Protein and solvent are put into a box with side length and periodic boundary conditions of 13.28 nm (for 6 × 2), 18.86 nm (for 12 × 2), and 13.31 nm (for 6 × 4), and electrostatic interactions are calculated by the particle mesh Ewald (PME) method ^20^. Stability of our structures is probed by following our molecular dynamics trajectories over 20ns, with only the last 10 ns used for analysis.

Most of our analysis is carried out within the tool set provided by GROMACS, with snapshots of configurations visualized by VMD ^21^. The distance between residues are defined as the average distance between heavy atoms in the sidechains of each residue; for example, the NZ atom on K16 and the OD1 atom on D23 are used to calculate the inter-chain salt-bridge between residues K16 and D23. We measure the solvent accessible surface area (SASA) by both the *g_sasa* and *POPS* ^22^ software tools, the later allowing one to separate the hydrophobic and hydrophilic areas. Since the two methods use different definitions of surface area, values may differ slightly. The binding energy are approximated with *MMPBSA.py* in AmberTools ^23^, with the setting igb=8 for the GBSA part.

### C. Aβ monomer and oligomer purification

Aβ monomers and oligomers were purified as described previously ^4^. Briefly, 0.5-1 mg of synthetic peptide was weighed into a sterile microcentrifuge tube and resuspended in 500 μL of 10 mM NaOH. After incubation at 25 °C for 30 min, the sample was loaded onto a Superdex-75 HR 10/30 size exclusion column preequilibrated in 20 mM Tris, pH 8.0 using either an AKTA FPLC (GE Healthcare) or BioLogic DuoFlow (BioRad) purification system. Fractions of 500 μL were collected at a constant flow rate of 0.5 mL/min. Aβ concentrations were determined using intrinsic tyrosine absorbance (ε = 1450 cm^−1^ M^−1^ at 276 nm) on a Cary 50 UV-Vis spectrometer (Agilent Technologies). To generate LFAOs, Aβ monomers (50-60 μM) were incubated with 5 mM C12:0 NEFA and 50 mM NaCl at 37 °C for 48 hours. LFAOs were then centrifuged at 20,000 *g* for 20 min before being purified via SEC as described above.

### D. ANS binding assay

ANS binding experiments were done as described previously ^2^. For experiments at varying pH, LFAOs were exchanged into 20 mM Tris at the appropriate pH using a 3.5 kDa molecular weight cut-off Slide-A-Lyzer MINI Dialysis Device (Thermo Fisher Scientific) following the manufactures protocol. Upon the addition of 100 μM ANS followed by 1 min of equilibration, fluorescence measurements of LFAOs (8, 6, 4, 2, 1, or 0.5 μM) were collected on a Cary Eclipse instrument (Agilent Technologies) by scanning the emission spectrum between 400 and 650 nm upon excitation at 388 nm. The area under the curve for each respective pH or NaCl titration experiment were then normalized from 0-1, and plotted as shown in the text. The data presented are representative of three independent experiments.

### E. Circular dichroism spectroscopy

Data were collected on a Jasco J-815 spectropolarimeter attached with a Peltier temperature controller. To a solution of LFAOs (1 or 8 μM), sodium dodecyl sulfate (SDS) was added to a final concentration of 1% (wt/v), followed by heating from 10-90 °C (ramp rate = 0.5 °C/min) while monitoring the signal at 206 nm every 1 min (hold time = 30 s, DIT = 32 s, bandwidth = 5 nm). The data was processed by normalizing from 0-1 (as shown in Fig 4a), followed by determining the apparent standard free energy using the Gibbs-Helmholtz equation (as shown in Fig 4b). The data presented are representative of three independent experiments.

### F. Atomic force microscopy

AFM samples were prepared following a previously published procedure ^3^. Freshly cleaved mica substrates were first treated with 150 μL of APTES (3-aminopropyltriethoxysilane) solution (500 μL in 50 mL of 1 mM acetic acid) for 20 minutes. The APTES solution was then decanted and rinsed three times with 150 μL DI H_2_O. The substrates were dried under a stream of N_2_ and stored in the desiccator for 1 hour. A 150 μL aliquot of Aβ solution (either 1 or 5 μM in 20 mM Tris-HCl, pH 8.0) was deposited onto the amine-treated mica substrates for 30 minutes to adsorb the proteins. The Aβ solution was then decanted and the samples were rinsed three times with 150 μL DI H_2_O. The samples were dried under a stream of N_2_ and stored in the desiccator until imaging.

AFM analysis of LFAOs was conducted using a Dimension Icon Atomic Force Microscope (Bruker) in PeakForce Tapping mode. AFM scanning was performed using NanoScope 8.15r3sr8 software and the images were analyzed in NanoScope Analysis 1.50 software. Imaging was performed using a sharp silicon nitride cantilever (SNL-C, nominal tip radius of 2 nm; nominal resonance frequency of 56 kHz; nominal spring constant of 0.24 N/m) in a standard probe holder under ambient conditions with 512×512 data point resolution.

## III. RESULTS

### I. LFAOs are two-layered rings

Detailed biophysical characteristics of LFAOs were obtained previously ^2-5, 7^. LFAOs display the presence of two aggregate distributions corresponding to sedimentation coefficients 5S and 7S (Fig 1a, reproduced from ^6^), corresponding to 12mers and more heavy assemblies of 12-24mers. The secondary structure determined by far-UV circular dichroism (CD) shows β-sheet structure that remains unchanged in an order of magnitude concentration difference (Fig 1b; reproduced from ^2^). However, two oligomer distributions were observed within the same concentration range on immunoblots (Fig 1b; *inset);* a band corresponding to 50-60 kDa (12mer) at 0.5 μM, and an additional band at 80-110 kDa (24mer) at 8 μM ^2^. Atomic force microscopy (AFM) analyses indicate spherical punctuate dot-like morphology for LFAOs with largely two distinct sizes corresponding to the sedimentation velocity analysis in panel “a” (arrows; Fig 2c). The solvated diameter determined from dynamic light scattering (DLS) ranged between 10-13 nm (Fig 2d). The concentration-dependent transition can be monitored by the increase in solvent exposed hydrophobic surfaces, as determined by 1-anilino naphthalene sulfonate (ANS) binding (Fig. 1e, •), and is consistent with an apparent dissociation constant (*K_d_*) of 0.1 μM ^2^. This transition between 12mer and 24mer LFAOs is absent in both monomer (☐) and fibril (Δ) control samples (Fig. 1e, reproduced from ^2^).

**Fig 1.**
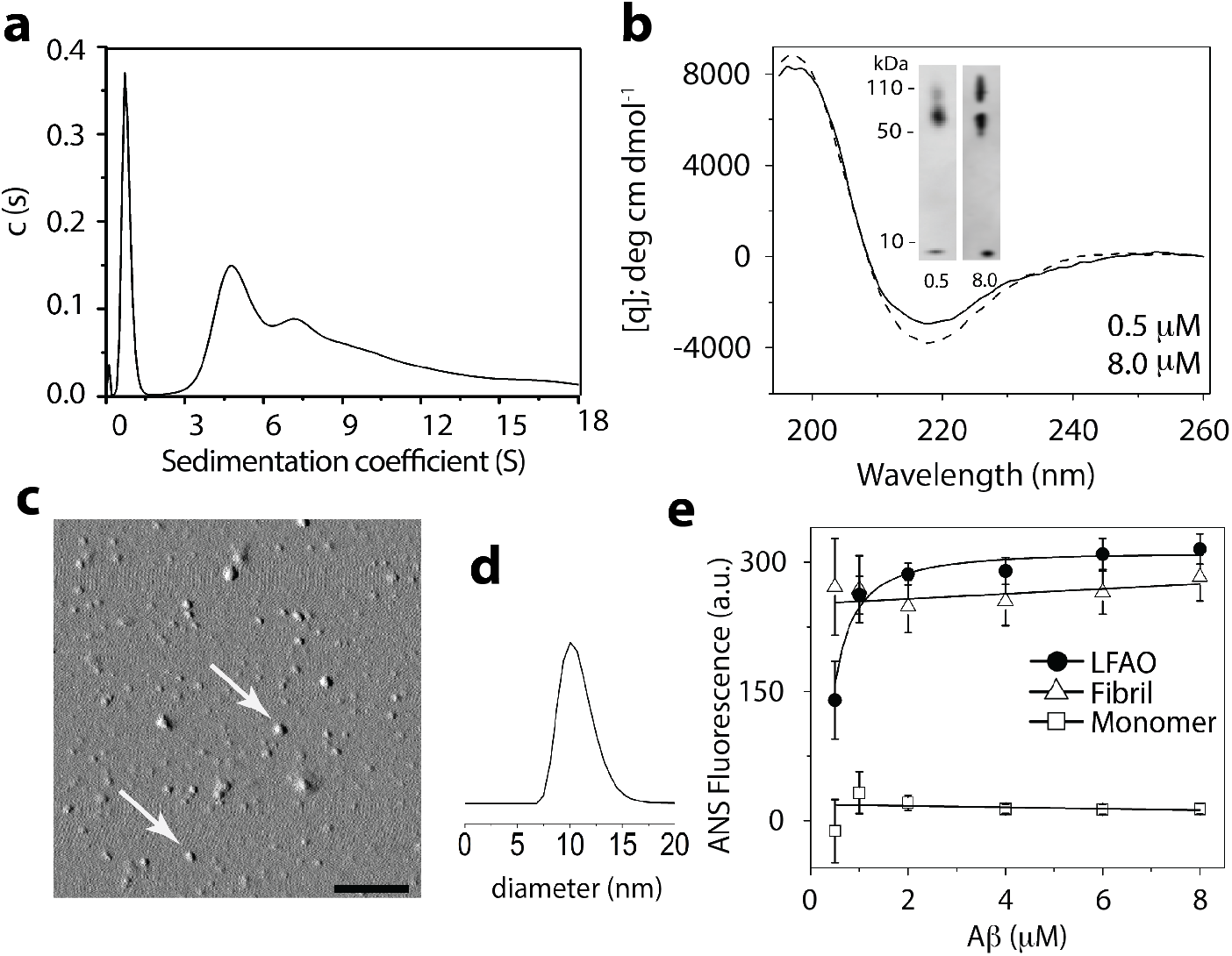
Biophysical characteristics of LFAOs (reproduced from the cited references). (a) LFAOs, as analyzed previously by analytical ultracentrifugation (5) and atomic force microscopy *(inset*, (3)). The single and double arrows represent LFAO 12mer and 24mer, respectively. (b) LFAO (●) concentration-dependent dynamics, as shown previously (2) using ANS fluorescence. Aβ monomers (□) and fibrils (Δ) are shown as controls. (c) Circular dichroism spectra and immunoblotting analysis *(inset)* of LFAOs at 0.5 (dashed) and 8 (solid) μM, as previously described (2). (d) AFM image shows punctuate spheres, scale bar represents 200 nm (reproduced from (3)). (e) DLS measurement shows a monodisperse species with diameter centered at 10 nm.

**Fig 2.**
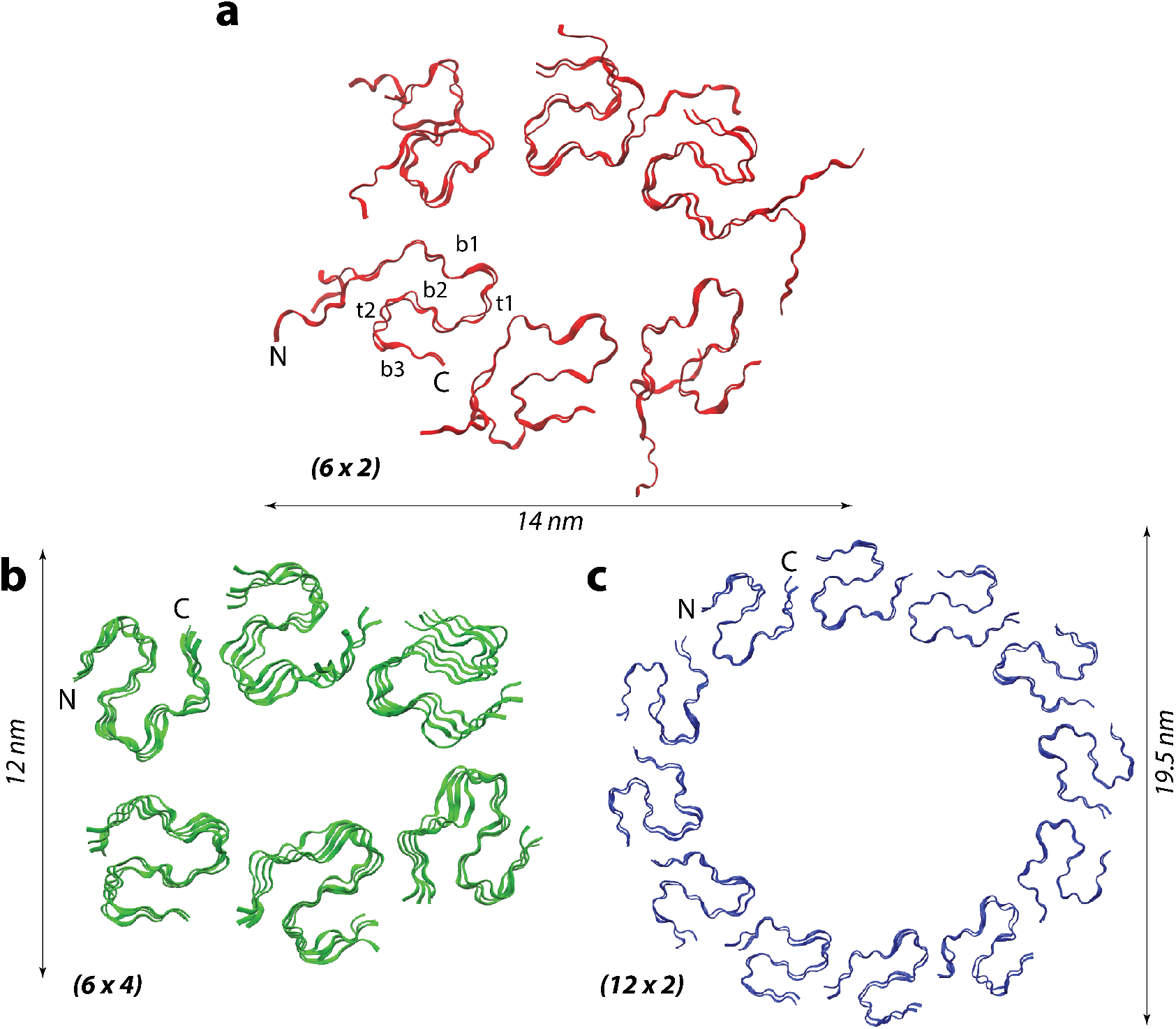
Models of LFAO 12 and 24mers. a) The backbone ribbon representation of our (6 × 2) model for LFAO 12mers built from two hexameric rings stacked on top of one another. β-sheets and turns are indicated as β and t, respectively. b and c) two possible models for LFAO 24mer assembly with (6 × 4) and (12 × 2), respectively.

**Fig 3.**
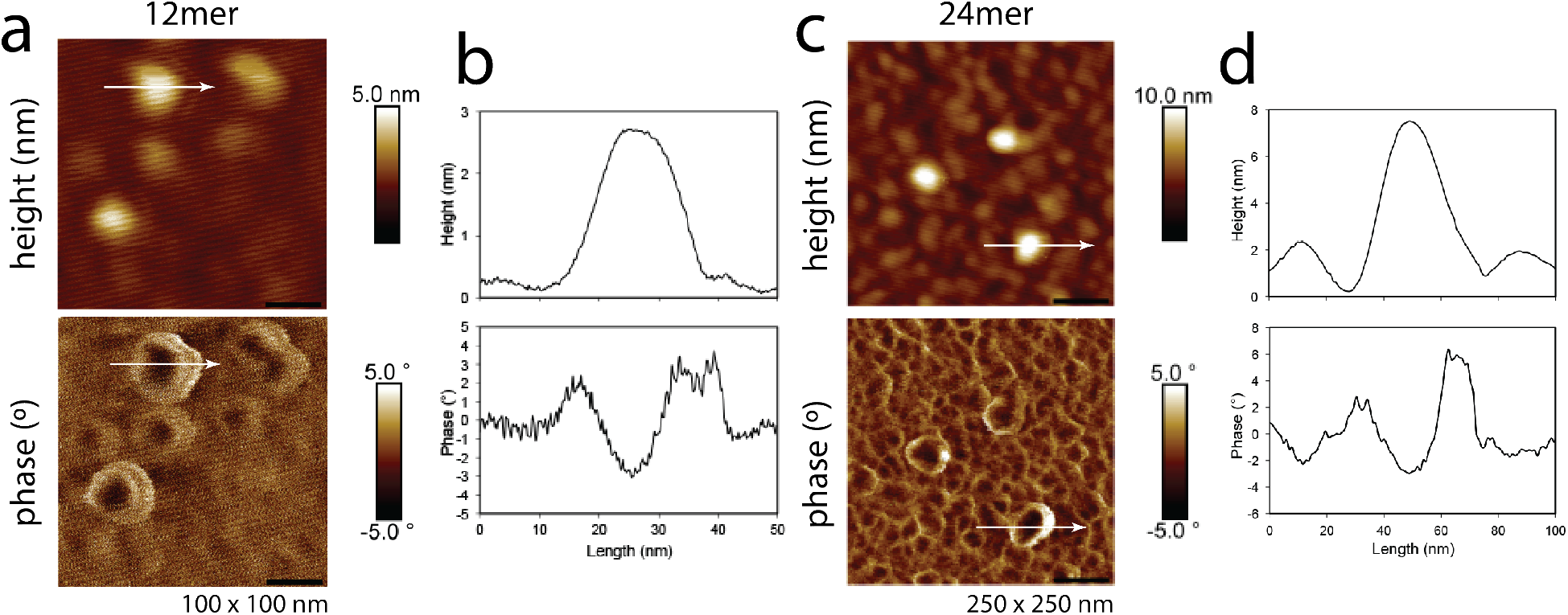
AFM morphology analysis of LFAO 12 and 24mers. a and b) Height and phase images of 1 μM LFAO samples (scale bar = 20 nm). Cross sectional analysis (XZ, shown in panel b) was conducted across the path indicated by the arrows. c and d) Height and phase images of 5 μM LFAO samples (scale bar = 50 nm). Cross sectional analysis (XZ, shown in panel d) was conducted across the path indicated by the arrows.

**Fig 4.**
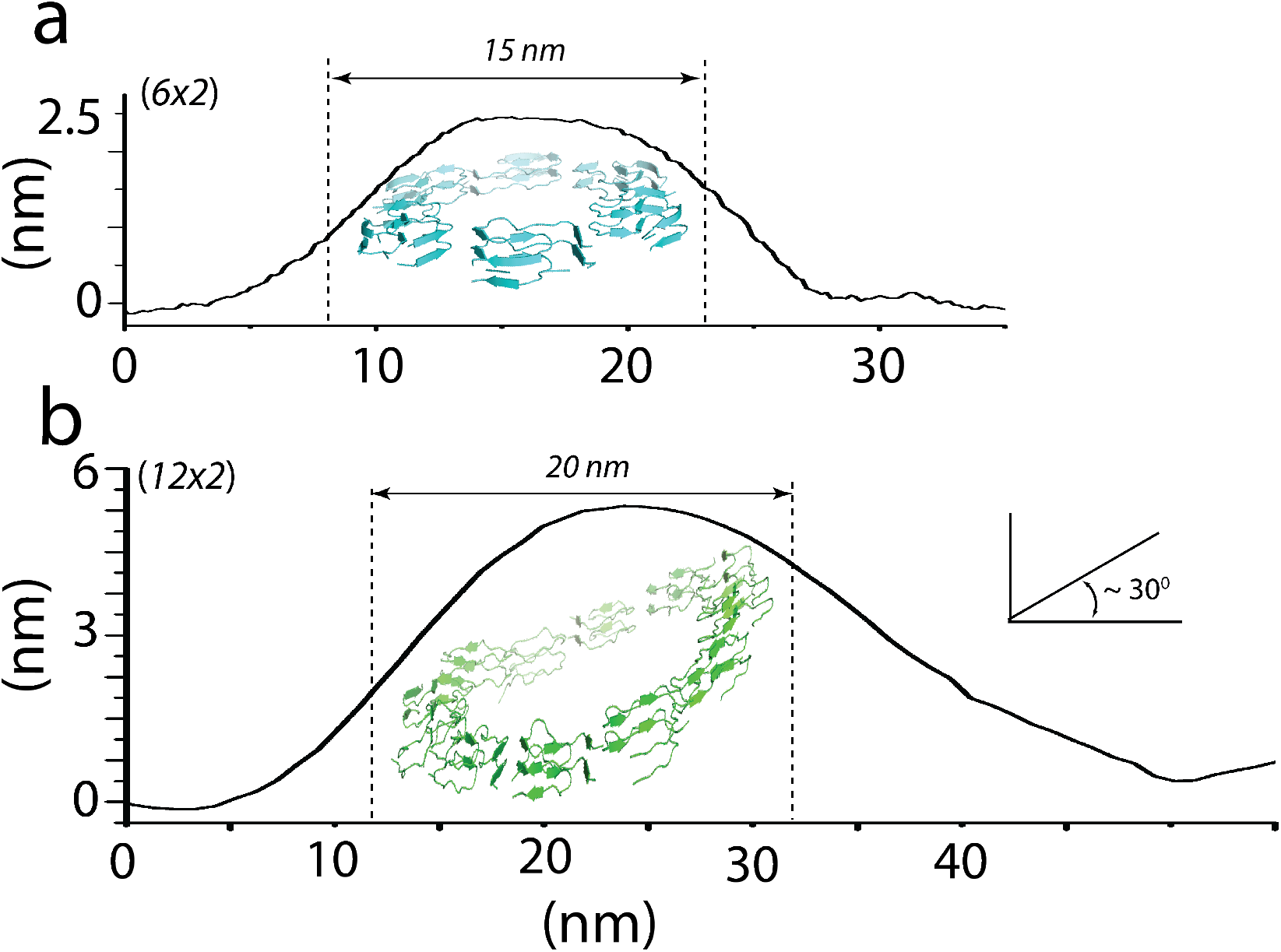
Experimental and simulation correlation for LFAO 12 and 24mers. a) The (6 × 2) structure is overlaid on to the height of smaller punctuate particles observed at low concentrations of LFAOs shown in Figure 3a and 3b. The height and diameter correlate with the observed ~ 3 and 14 nm, respectively for the (6 × 2) structure. b) The height of the larger spherical particles of LFAOs observed at higher concentrations in which a (12 × 2) structure is overlaid. The height of 5 nm can be accounted for only when the oligomer is adhered at a ~ 30° angle as shown. The corrected diameter of the species is also shown after accounting for the AFM tip radius.

In a recent paper ^24^, we have shown that unlike the more common, but less toxic Aβ40, Aβ42 polypeptide innately can form pore-like trimers and larger oligomers. This is because Aβ42 chains are able to assume a S-shaped three-stranded motif, while Aβ40 peptides are not stable in this form and instead take U-shaped conformations in fibrils ^15^. Building on our previously presented models ^24^ and guided by the experimental size measurements, we conjecture that the 12mer is organized as two ring-like hexamers stacked on top of one another (6 × 2) (Fig 2a) ^8^. In such an arrangement, the hydrated diameter of the oligomer measures 14-15 nm with a height of 3-4 nm, giving a flattened disc-like appearance. Note that oligomers with a similar structure have also been reported already earlier for Aβ peptide-fragments of various length ^25-27^. In order to model the heavier LFAO 24mers, two possible models were considered: a tetramer of hexamers (6 × 4), or a dimer of dodecamers (12 × 2). Both models are consistent with previous experimental results in their dimensions and size seen by AFM, which range between 14-20 nm of height observed ^2-3, 6^ (Fig 2b and 2c).

Our earlier AFM analysis on LFAOs showed a bimodal distribution of oligomers (Fig 1c), but we did not interrogate morphology of individual aggregates. In this study, we have employed a sharp silicon nitride cantilever with a lower resonant frequency and smaller spring constant to obtain higher resolution images. Height and phase images were obtained over small scan areas containing LFAOs of different diameters (Figure3). Height images indicate spherical particles with average diameters of 20 nm for the 12mers and average diameters of 40 nm for the 24mers. Average height for the 12mers is 2.5 – 3.5 nm and that of the 24mers is 5-8 nm. Phase images for both oligomers show a characteristic “donut” shape, indicative of differences in modulus and/or adhesion from the outer to the inner edge of the material, which we attribute to the existence of a cavity. Similar height and phase images have been reported for hollow nanoparticles. ^28^

To see the correspondence between the morphology obtained from AFM and the simulated models, the height analysis and various structural models were overlaid (Fig 4). A comparison of the dimensions derived from the AFM data for the 1 μM samples and those from (6 × 2) model shows a good agreement between the two (Fig 4a). Due to the large AFM tip diameter (2 nm) in relation to the size of the particles, the shape of the height plot was corrected to calculate the actual diameter of the oligomer, which yielded 15 nm for the 12mer as indicated in Fig 4a. The height and diameter obtained agree with the (6 × 2) model, which were 3 and 14 nm, respectively (Fig 2). Similarly, the heights of the larger spherical particles (24mer) obtained at higher concentrations (5 μM) were compared with the (12 × 2) model (Fig 4b). The corrected diameter (20 nm) corresponds to the one obtained from the model (19.5 nm). It is noteworthy that the slight increase in AFM heights (5-8 nm) in comparison to the height of the two layers in the 12 × 2 model (3-4 nm) can be accounted for when the oligomer is layered at ~ 30° angle on the mica surface (Fig 4b). On the other hand, the height measurements exclude the possibility of both a single-layered dodecamer ring (12 × 1) or a four-layered hexamer (6 × 4) structures. This observation parallels similar ones observed previously for Aβ42 oligomers ^27^. Note that our structural models are for residues 11-42 in the Aβ42 chains as residues 1-10 are flexible in all resolved fibril structures. To evaluate the effect of first 10 residues, we also constructed the full-length Aβ42 oligomers for (6 × 2) and (12 × 2) models with the N-terminal segment in the beginning in a random coil configuration that is allowed to relax in a molecular dynamics simulation over 10 ns. The orientation of the first ten residues stayed random, and their sole effect was that the diameter of (6 × 2) oligomers changed from 10.5 to 14.6 nm.

In order to understand why the heavier LFAO 24mers appear to form a dimer of dodecamers (12 × 2) instead of a tetramer of hexamers (6 × 4), we have simulated all three Aβ11-42 oligomer models, the (6 × 2) 12mer and the two 24mer models (6 x4) and (12 × 2), by atomistic molecular dynamics (MD). As we are neither modeling the association into the 12mers nor the transition between 12mers and 24mers, only relative short simulations are needed to explore energetics and stability of these models. Note that the experimental measurements were obtained at a pH=8, i.e., under neutral/alkaline conditions, which were modeled in our simulations by changing the H13 and H14 residues into a deprotonated state (named by us the HIE state). Each system is followed in two independent runs. In Table 1 we list the solvent accessible surface area (SASA) and the binding energy as approximated by the MMGBSA approach. This approximation is justified because we are not interested in absolute values for the binding energies of the three models but only in the qualitative differences between them.

**Table I.** The solvent accessible surface area (SASA) and the binding energy of the three considered oligomer models under neutral pH conditions. Shown are for all quantities the averages (Avg) as obtained from two runs of 20 ns and their standard deviations (StD). The binding energy was calculated in a MMGBSA approximation and is composed by four terms: the van der Waals energy E_vdW_, the electrostatic energy E_elect_, a generalized Born approximation of the solvation energy E_GB_, and solvent surface tension interaction term E_surf_. The SASA values are calculated by two different tools: g_sasa and POPS. In POPS, the SASA values can be further separated into hydrophobic and hydrophilic contributions. The binding energy are defined as follows: Δ(A)=E_(12×2)-2×E_(6×2), Δ(B)=E_(6×4)-2×E_(6×2)).

| Binding energy / kcal/mol |  | (6 x 2) |  | (6 x 4) |  | (12 x 2) |  |
| --- | --- | --- | --- | --- | --- | --- | --- |
|  |  | Avg. | StD | Avg. | StD | Avg. | StD |
|  | E<sub>VDW</sub> | -690.4 | 2.2 | -953.1 | 0.2 | -1445.9 | 1.0 |
|  | E<sub>elect</sub> | -213.3 | 4.0 | 478.0 | 46.8 | -429.3 | 57.3 |
|  | E<sub>GB</sub> | 371.9 | 1.5 | -259.3 | 51.6 | 757.5 | 52.7 |
|  | E<sub>surf</sub> | -89.5 | 0.1 | -113.6 | 0.5 | -186.8 | 1.3 |
|  | Total | -621.3 | 0.4 | -848.0 | 4.4 | -1304.5 | 6.9 |
| SASA / Å<sup>2</sup> | g_sasa | 250.4 | 1.3 | 339.1 | 0.3 | 495.2 | 3.0 |
|  | POPS: All | 258.4 | 1.3 | 400.7 | 0.6 | 524.0 | 3.2 |
|  | Hydrophobic | 163.9 | 0.4 | 255.2 | 0.4 | 332.2 | 2.8 |
|  | Hydrophilic | 94.5 | 0.4 | 145.4 | 0.2 | 191.8 | 0.3 |
|  | Delta of binding energy kcal/mol |  |  | Avg |  | StD |  |
| | $\Delta(A)$ | | | -58.04 | | 3.9 | |
| | $\Delta(B)$ | | | 392.80 | | 1.8 | |

Comparing the SASA of the three models we note that the values for the (12 × 2) model of the 24mers is about double that of the (6 × 2) 12mer, while the corresponding SASA of the (6 × 4) 24mer model is less than 1.5 times that of the 12mer. Hence, based on the ANS binding data, which showed that the solvent accessible hydrophobic surface area doubles from LFAO 12mers to 24mers (Fig 1d), we conclude that the (12 × 2) model is the more likely structure for the 24mers. This model has a dimeter of 15.7 nm, which extends to 19.5 nm for the full-sized model (including the first ten N-terminal residues), also in agreement with the experimentally measured dimensions.

The (12 × 2) model for the 24mer is also favored by the binding energies shown in Table 1. The (12 × 2) structure has binding energies more favorable than the binding energies of two (6 × 2) models (see the difference Δ(A) = E_(12×2)-2×E_(6×2) in Table 1); while the binding energy of the 6 × 4 model is substantially higher than that of two isolated 6 × 2 models (see the difference Δ(B) = E_(6×4)-2×E_(6×2) in Table 1), disadvantaging formation of this assembly. In order to show why the 12 × 2 model is more favorable than the (6 × 4) model we neglect entropic contributions and approximate the binding energy of a 12mer by;

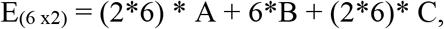

where A is interaction between chains *within* a ring, B the interaction between chains of *neighboring* rings, and C the interaction of chains with *surrounding water* (i.e. proportional to exposed surface). With the same definitions, one finds that the binding energy of a (12 × 2) would be;

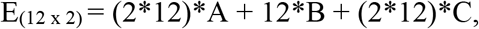

i.e, Δ(A)=E_(12×2)-2×E_(6×2) = 0. On the other hand, the binding energy of (6 × 4) would be;

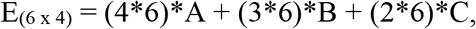

i.e, Δ(B)=E_(6×4)-2×E_(6×2)) = 6*B – (2*6)*C. This difference describes in first approximation how much more unfavorable the (6 × 4) model is over the (12 × 2) model, and results from repulsive interactions between the (6 × 2) 12mers. The histidine side chains at positions 13 and 14 orient themselves towards one side of the hexamer ring in the upper layer, while similar histidines on the congruent Aβ42 unit on the lower layer hexamer ring do so on the opposite side. Hence, the positioning of the histidine chains hinders further association of hexamer rings (leading to steric clashes), which would make a (6 × 4) structure thermodynamically expensive. The above results suggest that the transition between 12mers and 24mers is not a simple stacking of two 12mers. As the repulsive interaction between (6 × 2) 12mer prohibits stacking and leads to a large unfavorable binding energy, a reorganization of chains is needed, leading to a (12 × 2) structure where the histidines (H13 and H14) are moved ~ 95° perpendicular to the axis of the oligomer, thus preventing potential charge repulsion due to protonation. Furthermore, this transition also results in the exposure of hydrophobic residues along either side of oligomer face, see Table 1, evident from increase in ANS binding (Fig 1d).

This scenario is also consistent with the observation that the transition between 12mer and 24mer depends on concentration, see (Fig 1b and 1c) ^2^, and can be understood from the energetics of our (6 × 2) 12mer and (12 × 2) 24mer models. At low concentrations, the 12mers are separated and experience little repulsive forces. At the same time, the gain in energy from forming a 24mer is negligible, and therefore the equilibrium between the two forms shifts towards the 12mer. On the other hand, above a critical threshold, the distance between (6 × 2) 12mer rings becomes so close that the repulsive interaction between them becomes noticeable. At that point it becomes energetically more favorable to rearrange and form (12 × 2) assemblies, i.e., the equilibrium is shifted toward the 24mers.

## II. Dynamics based on stability

While the above reasoning is plausible, more evidence is needed to support our models for 12mer and 24mer, and for the implied mechanism of the concentration-dependent transition between the two forms. Since the transition from 12 to 24mer involves exposure of charged histidines (Fig 1d), we reason that the transition is caused by electrostatic interactions. In an effort to obtain molecular details on such a possibility, we investigated the effect of buffer pH and salinity on 12 to 24mer transition using the ANS binding assay. This transition was previously established at pH 8.0 in low ionic strength conditions (absence of salt) (□; Fig 5a and 5b). Upon decreasing the pH from 8.0 to 5.0, the 12 to 24mer transition was less pronounced with a decrease in binding affinity (Fig 5a). Similarly, systematically increasing the ionic strength also resulted in a diminished ability of 12mers to convert to 24mers (Fig 5b). The decrease in pH near the isoelectric point (pI) of Aβ (5.5) resulted in a reduction in propensity of 12- to 24mer conversion. This indicates the direct involvement of favorable electrostatic interactions, as the abrogation of charges (near the pI) diminishes the 12 to 24mer conversion (Fig 5a). To further ascertain the role of the protonated of H13 and H14 residues in dimerization, we generated H13A/H14A double mutant of Ab42 and generated oligomers in similar conditions as that of LFAOs (data not shown). The dimerization of isolated oligomers was then monitored by ANS binding at pH 5, where LFAOs showed weakest 12 to 24mer transition. Upon titration, we observed that the double mutant specifically rescued the dimerization of the oligomers at pH 5.0 where the histidines would be protonated (Fig 5c). Specifically, this consolidates the idea derived from the structures regarding the involvement of protonation/deprotonation events in such a transition. To further investigate differences in 12mers and 24mers, thermodynamic stability analysis was performed in the presence of a denaturant (sodium dodecyl sulfate, SDS). In these experiments, the conversion of β-sheet structure adopted by LFAOs to an a-helix in the presence of SDS was monitored by far-UV CD at 206 nm as a function of temperature (Fig 6a). The equilibrium data were processed (as described in the experimental) to obtain the apparent Gibbs free energy at 37 °C, which were found to be −0.879 and −0. 489 kcal/mol for LFAO 12mer (1 μM) and 24mer (8 μM), respectively (Fig 6b). This difference of 0.4 kcal/mol is minimal and insignificant, revealing that both isoforms have similar thermodynamic stability.

**Fig 5.**
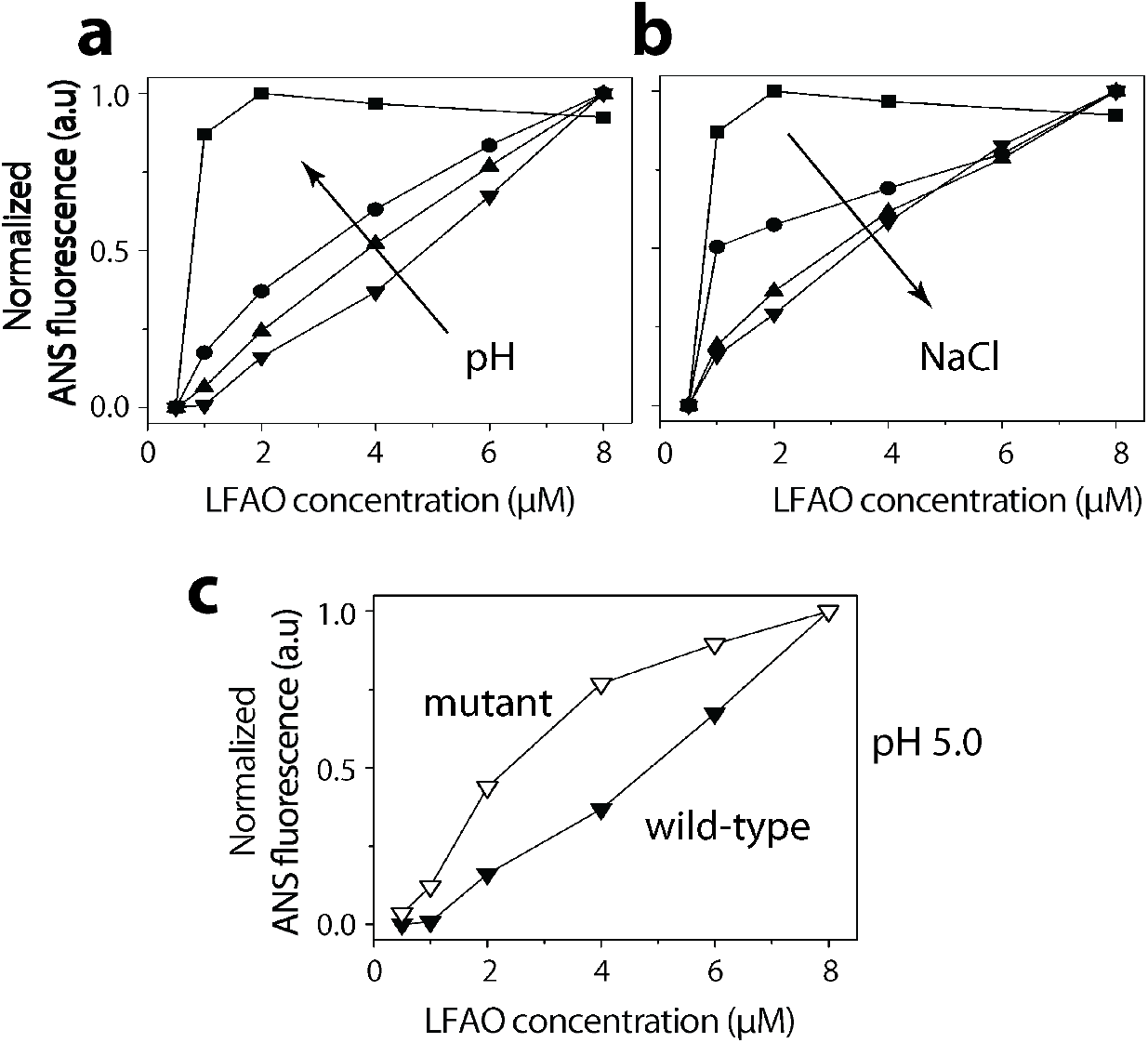
LFAO conformational dynamics at varying NaCl and pH. Normalized ANS fluorescence for 0.5, 1, 2, 4, 6, and 8 μM LFAOs at varying (a) NaCl (0, ■; 50, ●; 100, ▲; or 200 mM, ▼); (b) pH (8, ■; 7, ●; 6, ▲; or 5, ▼) respectively, and c) rescue of dimerization at pH 5 by H13A-H14A double mutant. Data were processed as described in the Experimental section.

**Fig 6.**
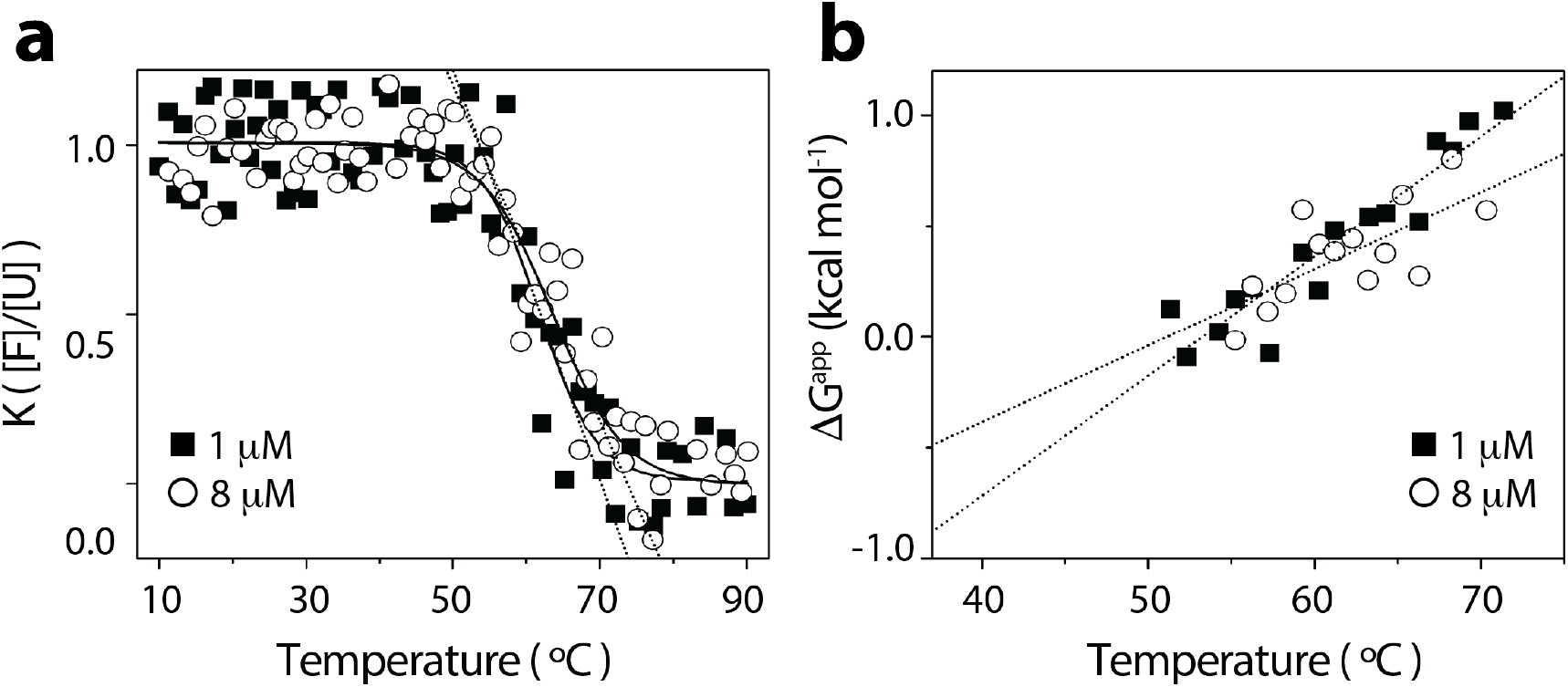
Thermal stability analysis of LFAO 12mer and 12-24mers. (a) Normalized circular dichroism spectra, collected at 206 nm, of 1 (■) and 8 (O) μM LFAOs upon the addition of SDS (1%) followed by heating to 90 °C. (b) The equilibrium constant (K, [folded]/[unfolded]) from panel (a) was used to determine the apparent standard free energy change (ΔG_*app*_) for both 1 (■) and 8 (O) μM LFAOs, which was extrapolated to be −0.879 and −0. 489 kcal/mol at 37 °C (y-intercept), respectively. The data were process as described in the Experimental section.

In our simulations, we model the switch from neutral to acidic conditions by repeating our simulations of the three systems (the (6 × 2) for the 12mers, and the (6 x4) and (12 × 2) model for the 24mers), with protonated histidine 13 and 14, i.e., setting the charge state to positive (called HIP in this paper). All other parameters in the two sets of simulations are identical. Data for the pH<7 simulations are summarized in Table 2. Comparing binding energies between charged (pH < 7) and neutral forms, it appears that the energy differences between 24mer (12 × 2) and 12mer (6 × 2) favor the 24mers in the charged state more than in the neutral state ((ΔΔG) = ~ 13 kcal/mol). This is unexpected as the experimental data shows a faster transition toward 24 mer at pH ≥ 7. Hence, the sharper transition between 12mer and 24mer is *not* because at pH ≥ 7 the 24mer is energetically more favored over the 12mer than in acidic conditions. Instead, the sharper transition at pH ≥ 7 is because the repulsion between 6 × 2 dodecamers is much higher for the charged forms than the neutral forms (Table 1 and 2). In other words, the repulsion between the 12 mers is larger at low pH values than in the neutral range where a faster 12 to 24mer transition was observed. Hence, we conjecture that the less pronounced transition at low pH is because larger concentrations are needed to overcome the stronger repulsion between the 12mer than at neutral or higher pH. Note also that while the solvent accessible surface area does not differ between the charged and neutral forms in the (6 × 2) model, a difference in SASA is observed for the (12 × 2) structure. For neutral conditions (HIE), the (12 × 2) structure of a 24mer exposes roughly two times (332 Å^2^; Table 1) the hydrophobic suraces of the (6 × 2) 12mer (164 Å^2^; Table 1), an observation that the experimental results concurr with (Fig 1b). On the other hand, under acidic conditions (HIP), the 24mer exposes with 347 Å^2^ (Table 2) more than double the hydrophobic surface of two 12mers (164 Å^2^; Table 2), making formation of 24mer less favorable), which further agrees with the experimental observations.

**Table II.** The solvent accessible surface area (SASA) and the binding energy of the three considered oligomer models under neutral pH conditions. Shown are for all quantities the averages (Avg) as obtained from two runs of 20 ns and their standard deviations (StD). The binding energy was calculated in a MMGBSA approximation and is composed by four terms: the van der Waals energy E_vdW_, the electrostatic energy E_elect_, a generalized Born approximation of the solvation energy E_GB_, and solvent surface tension interaction term E_surf_. The SASA values are calculated by two different tools: g_sasa and POPS. In POPS, the SASA values can be further separated into hydrophobic and hydrophilic contributions. The binding energy are as follows: Δ(A)=E_(12×2)-2×E_(6×2), Δ(B)=E_(6 × 4)-2 × E_(6 × 2)).

| Binding energy / kcal/mol |  | (6 x 2) |  | (6 x 4) |  | (12 x 2) |  |
| --- | --- | --- | --- | --- | --- | --- | --- |
|  |  | Avg. | STD | Avg. | STD | Avg. | STD |
|  | E<sub>VDW</sub> | -696.1 | 5.2 | -903.6 | 6.3 | -1493.4 | 20.7 |
|  | E<sub>elect</sub> | -15.4 | 81.7 | 972.9 | 95.8 | 130.3 | 65.2 |
|  | E<sub>GB</sub> | 167.1 | 73.0 | -760.9 | 87.0 | 212.6 | 48.8 |
|  | E<sub>surf</sub> | -91.0 | 0.1 | -108.3 | 0.5 | -197.6 | 1.7 |
|  | Total | -635.3 | 3.4 | -800 | 15.6 | -1348.0 | 6.1 |
| SASA / Å<sup>2</sup> | g_sasa | 249.4 | 1.6 | 331.9 | 1.1 | 511.3 | 2.3 |
|  | POPS: All | 257.6 | 1.8 | 389.2 | 0.9 | 542.4 | 2.9 |
|  | Hydrophobic | 163.6 | 0.6 | 249.5 | 0.6 | 346.9 | 2.2 |
|  | Hydrophilic | 94.0 | 0.5 | 139.7 | 0.4 | 195.5 | 0.7 |
|  | Difference in binding energy kcal/mol |  |  | Average |  | STD |  |
| | $\Delta(A)$ | | | -71.0 | | 6.4 | |
| | $\Delta(B)$ | | | 466.2 | | 4.4 | |

## IV. DISCUSSION

What is the cause for the above described changes in the way the 12mer and 24mer differ at low pH and neutral pH, changes that in turn modulate the concentration-dependent transition between 12mer and 24mer? These changes have to be connected with differences in the two geometries, and with the way these differences are modulated by pH. One possibility is the way residues are exposed to the solvent. Fig. 5 shows the per-residue differences in solvent exposed surface area between HIE and HIP states for (6 × 2) and (12 × 2) structures. While on average the SASA does not differ between charged and neutral forms in (6 × 2) 12mer, the turn region t1 between β1 and β2 (see Fig 2) is more exposed under neutral conditions (HIE) than under acidic conditions (HIP), while most other residues are more exposed in the HIP state than in the HIE state. A similar picture is seen for the (12 × 2) 24mer, only that here the difference in exposure of residues in the turn region is much smaller than for the 12mer; while the overall exposure of surface to the solvent is larger under acidic conditions (HIP) than under neutral conditions (HIE). Hence, our solvent accessible surface differences indicate that the pH-modulation of the transition between 12mer and 24mer involves this turn region.

This observation is confirmed by Fig 7 where the per-residue contributions to the binding energy are shown. The contribution of binding energy for each residue was calculated in three different ways: first, the difference between HIE and HIP states for (6 × 2) and (12 × 2) models (Fig 7a), second, the difference in binding energies between two times that of (6 × 2) model and a (12 × 2) model (Fig 7b). In both cases no apparent signal is seen in the figures. However, when looking into the difference in binding energy of two isolated (6 × 2) models minus the binding energy of the (6 × 4) model (Fig 7c), which is a measure for the maximal repulsion between two (6 × 2) models, a clear signal is observed. The only segment where there is a difference between HIE and HIP are residues 20 to 28, which include the turn region between β1 and β2, and other residues located on packing surface that directly interact with residues on the neighboring fold. For this segment, the binding energy contribution is for neutral pH similar between two isolated (6 × 2) rings and when fully associated as a (6 × 4) assembly, while under acidic conditions the binding energy contribution from these segments favor isolated (6 × 2) assemblies. This is consistent with Fig 5a which shows that under acidic conditions the residues in this segment are less exposed to solvent than under neutral conditions.

**Fig 7.**
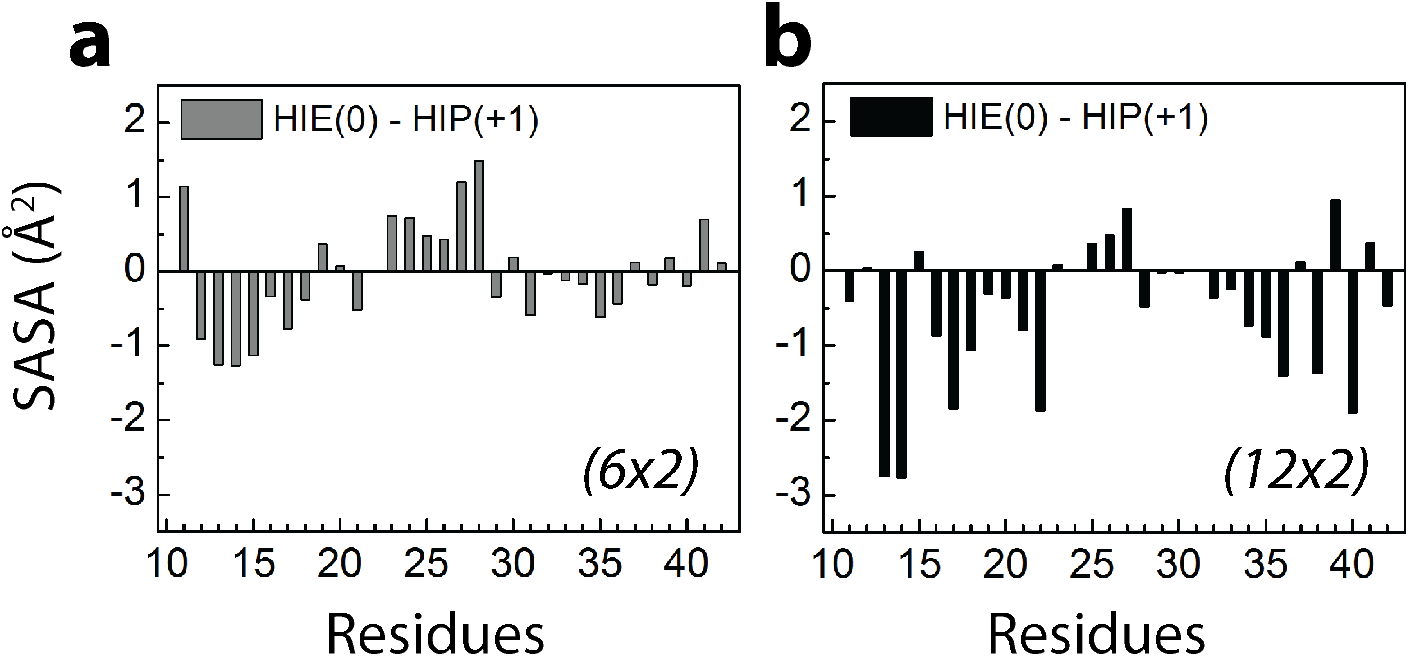
Difference in the solvent accessble surface area (SASA) contributions of single residues between acidic conditions (HIP) and neutral conditions (HIE) for the two Aβ42 oligomer models: (a) the (6 × 2) 12mer and (b) the (12 × 2) 24mer.

Visual inspection of this segment in the (6 × 2) model for both HIE and HIP systems shows that this region is slightly more distorted in HIP than in HIE (Fig 8). This distortion is related with (and can be quantified by) a weakening of the salt-bridge between K16 and E22 or D23 (mainly the K16-D23, see ^29^) that is formed between neighboring chains. In order to demonstrate this point, we have calculated the average distance of all corresponding salt-bridge pairs between the O^δ1^/O^δ2^ atoms on D23 and Nζ atom on K16, and the distributions of such distances for the HIP and HIE models are shown in Fig 10.

**Fig 8.**
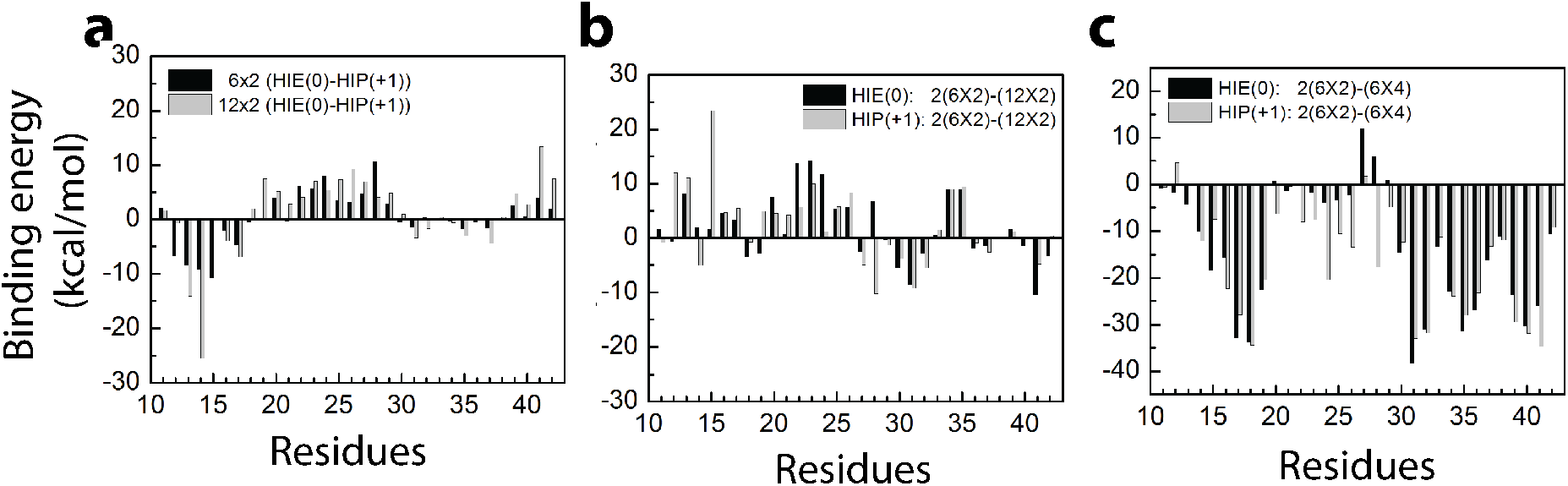
Differences in the binding energy contributions of each residue in the various models. (a) Difference between the values measured under acidic conditions (HIP) and neutral conditions (HIE) for both (6 x2) 12mer and (12 x2) 24mer; (b) difference between two times the value measured for the (6 × 2) 12mer and the value measured for the (12 x2) 24mer, data are for both acidic conditions (HIP) and neutral conditions (HIE); (c) difference between two times the value measured for the (6 × 2) 12mer and the value measured for the alternative (6 x4) model for a 24mer, data are for both acidic conditions (HIP) and neutral conditions (HIE).

Compared with the neutral state (HIE), the acid state decreases the stability of the inter-chain salt-bridge in the (6 × 2) model. How does the different charge states of the histidine lead to this effect? In HIP, residues 13 and 16 are both positively charged and the repulsive interaction between them distorts the geometry of the salt-bridge between residue K16 and either D23 or E22. In order to quantify the repulsive interaction between the two positively charged H13 and K16, we have calculated the average distance between the mass center of imidazole on H13 and Nζ atoms on K16 on the same chain and drawn the distribution of distances between H13 and K16 in Fig 10. The geometry in the 24mer (a two-layer dodecamer ring) is such that the different charge states of H13 does not change the average distance between H13 and K16, and therefore also does not weaken the salt bridge K16-D23 (E22), see Fig. 9. On the other hand, for the 12mer (a two-layer hexamer ring) the distribution of distances between residues H13 and K16 is shifted towards larger values for acidic conditions reflecting the repulsive interaction between the two residues under these conditions. This is not seen for the 24mer because for a (12 × 2) structure, the same histidines (H13 and H14) are moved ~ 95° perpendicular to the axis of oligomer, and thus preventing potential charge repulsion due to protonation. The net-effect of this repulsive interaction between the charged histidine and the lysine K16 is a weakening of the salt bridge K16-D23 (E22), (Fig. 10), which in turn reduces the stability of the turn region between the β1 and β2 strands and the hydrophobic core region for the peptides in the 12mer under acidic conditions. The binding energies (Fig. 8c) indicate that this distortion leads to a larger repulsive interaction between (6 × 2) structure, shifting the equilibrium toward the 12mer.

**Fig 9.**
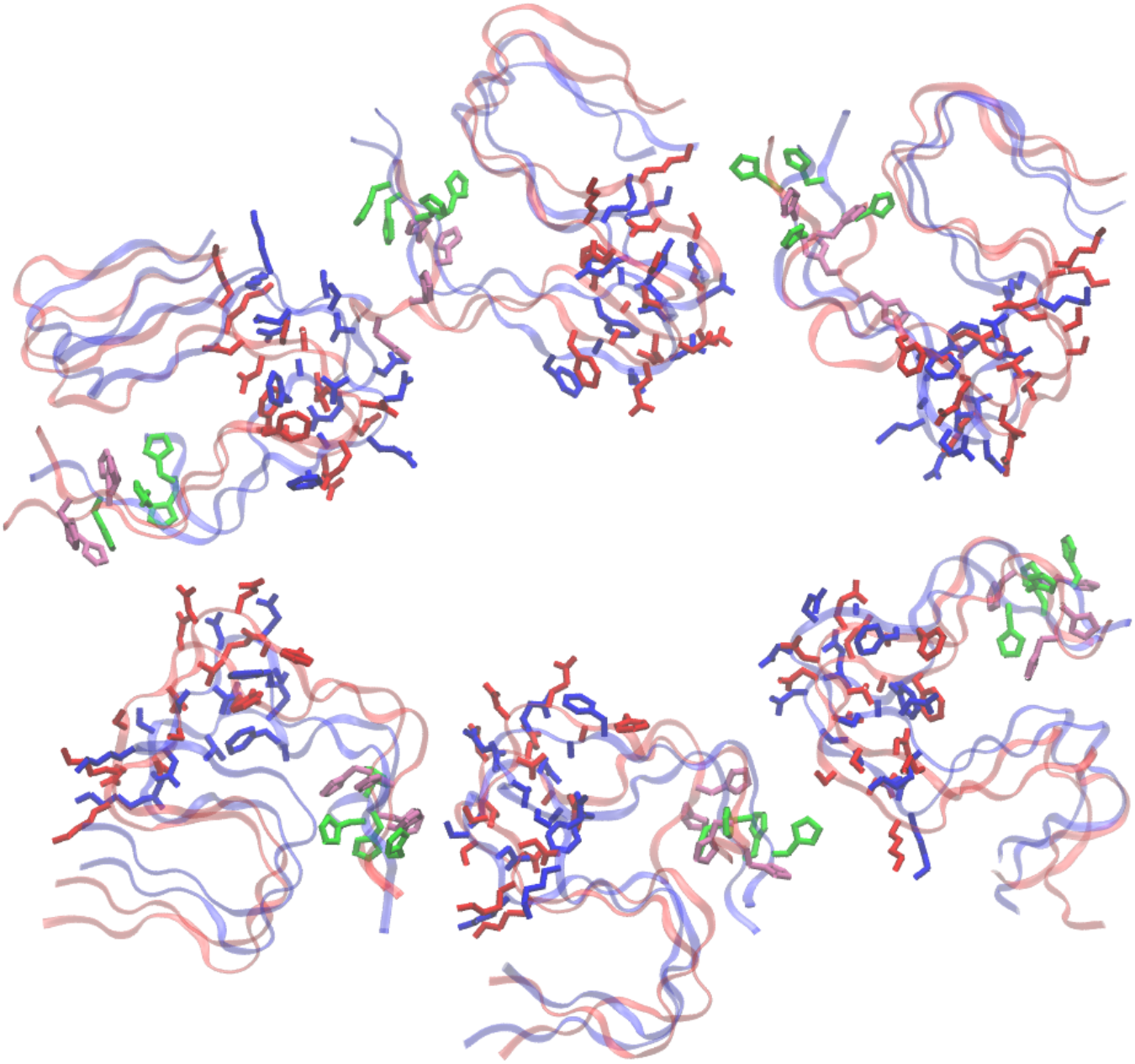
Final structures of the (6 × 2) model for neutral pH (HIE, blue) and acidic conditions (HIP, red) states after a 20ns molecular dynamics trajectory. The sidechains of residues 20 to 28, where the contributions to binding energy differed mostly with pH, are shown in bond representation. The histidine residues H13/H14 are colored in green (HIE) and mauve (HIP).

**Fig 10.**
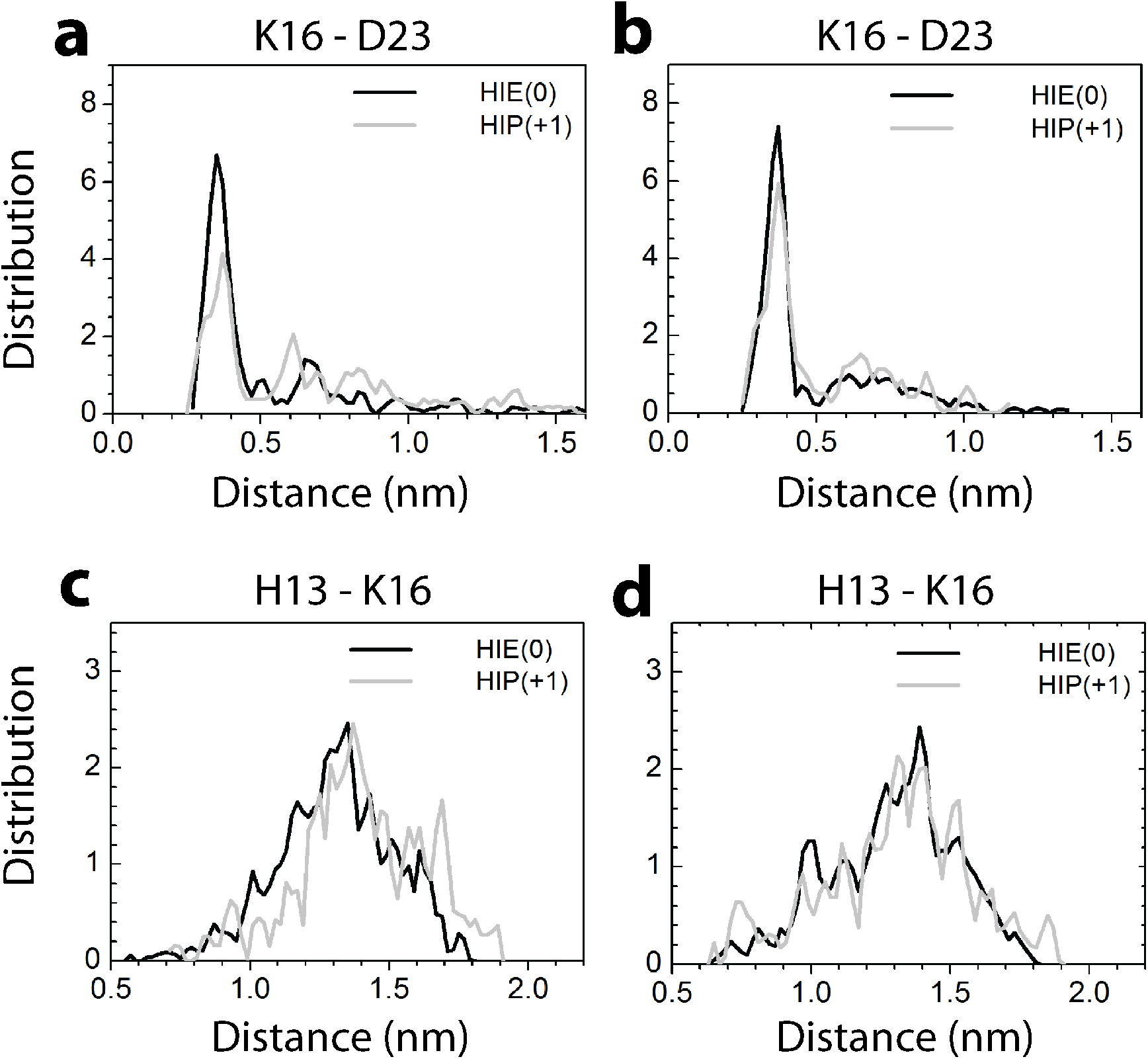
Distance distribution plot. The distribution of distances between the inter chain salt-bridge forming residue 16 (a positively charged lysine) K16 and residue 23 (a negatively charged aspartic acid) D23 in the (6 x2) 12mer (a) and the (12 x2) 24mer. The corresponding distribution of distances between residue 13 (histidine) H13 and residue 16 (a positively charged lysine) are shown in (c) for the 12mer and in (d) for the 24mer.

## V. CONCLUSIONS

Combining a variety of biophysical measurements and molecular dynamics simulations we put forward models for the fatty acid catalyzed Aβ42 assemblies called LFAOs. The experimentally observed existence of 12 and 24mer LFAOs in a concentration-dependent manner agrees with the models generated by simulations. First, ring-like two-layered assemblies were derived for both 12 and 24mers (two stacked hexamer rings in the case of the 12mer, and two stacked dodecamer rings in the case of the 24mer). The diameter and height of the two oligomers agree well with their morphology in AFM images that appear as punctuate spherical particles. More importantly, the most noticeable feature of the oligomer structure, the ring structure with a cavity in the middle, is also in agreement with the phase changes observed in AFM images. Furthermore, the determination that protonation of histidine side chains (H13 and H14) to be the key in determining the structural conversion of (6 × 2) to (12 × 2) is also supported by the experiments on both wild-type and double alanine mutant of the histidines. Finally, the energetic contributions calculated from the models are also in agreement with the free energy changes observes experimentally. Together, both simulations and experiments point out to the proposed models for the LFAO structure and dynamics that remained elusive thus far.

It is noteworthy that the disc-like oligomers of LFAOs observed here have also been observed for Aβ by other groups ^25-27^. Based on AFM imaging, ring-like, spherical low molecular oligomers are observed to be transiently formed before the formation of high molecular weight oligomers, which then laterally associate to form protofibrils ^25^. Economou and colleagues observed that even at low concentrations, Aβ42 but not Aβ40, form ring-like hexamers that convert to dodecamers, which consequently seed protofibril formation ^26^. Our observations on LFAO structure and propagation support our previous reports and add much needed detail. The studies undertaken in our labs have confirmed that LFAOs are ring-like dodecamers at low concentrations ^6, 8^. Investigations on the mechanism of LFAO propagation suggested that LFAO 24mers are formed at higher concentrations, which grow larger to form a key intermediate on route to fibril formation in a three-step mechanism ^4^. The (12 × 2) structure observed for 24mers explains the fact that 24mers, and not (6 × 2) 12mers, are able to faithfully propagate by associating with one another mediated by increased hydrophobic surface interactions with 24mer units of LFAOs as observed previously ^25, 27^. Overall, these observations are in agreement with those described previously.

Perhaps the intriguing and enigmatic properties of LFAOs are; (i) the ability of 12mers to self-replicate in the presence of monomers, (ii) to convert to 24mers in a concentration dependent manner and, (iii) the striking differences in the pathogenicity of 12 and 24mers. While both oligomeric forms are pathogenic, LFAO 12mers are more apoptotic to neuroblastoma cells than the 24mers ^2^. LFAOs also induce acute CAA in transgenic mice brains selectively, although it remains unclear which form of LFAO is responsible for this phenotype ^3^. The results presented here bring out the molecular signatures that are responsible for the structural and functional differences between LFAO 12 and 24mers and provide insights into the structure and mechanism by which LFAOs behave and become neurotoxic. At elevated concentrations, (6 × 2) LFAOs form 24mers by adopting a (12 × 2) structure accompanied by reorganization in the assembly, which exposes the hydrophobic residues along either side of oligomer face, an observation also supported by an increase in ANS binding. Such a reorganization increases the susceptibility of 24mers for further oligomer associations mediated largely by hydrophobic interactions. This is indeed supported by the fact that LFAO 24mers are able to propagate morphologically distinct fibrils made by repeats of LFAO units ^4^.

Insights into the molecular underpinnings of oligomer behavior is much needed to understand AD pathology and for future therapeutic interventions. This work is a step toward advancing our knowledge into this critical area, providing insights into the structure and mechanism by which LFAOs behave and become neurotoxic.

## AUTHOR CONTRIBUTIONS

VR and UHEH conceptualized and conducted the research. DND performed biophysical experiments and data analysis while WX conducted molecular dynamics simulations and analyzed the results. KS and SEM performed the AFM analysis. All authors were involved in manuscript writing and editing.

## ACKNOWLEDGEMENTS

The authors would like to thank the following agencies for financial support: National Institute of General Medical Sciences (R01GM120634, to UH and VR), NSF Graduate Research Fellowship Program (NSF 1445151) (to DND), NSF NRT 1449999, and NIH R15GM123431-01 (KAS and SEM). Simulations were done on the SCHOONER cluster of the University of Oklahoma. The authors declare no conflicts of interest. The authors also thank Ms. Jhinuk Saha and Ms. Morgan Malone for their help with DLS and double mutant experiments.

